# A biologically supported global RNA architecture of cucumber mosaic virus satellite RNA

**DOI:** 10.64898/2026.08.21.746196

**Authors:** Na Li, Yingying Gao, Siyuan Ren, Zhouhang Gu, Dongliang Yu, Qiansheng Liao, Zhiyou Du

## Abstract

Satellite RNAs (satRNAs) are parasitic subviral agents whose biological activities are mediated largely by specific sequence determinants and structured RNA elements. Establishing biologically supported global RNA structures is therefore essential for understanding how RNA architecture underlies satRNA functions. Cucumber mosaic virus (CMV) satRNA is one of the best-characterized models for investigating satRNA structure-function relationships; however, a biologically supported global RNA architecture of CMV satRNA has yet to be established. Here, we applied AlphaFold3 modeling to predict the global structure of CMV satRNA T1 (sat-T1). Initial full-length structure modeling generated multiple long-distance interactions that lacked biological support. We therefore used fragment-based modeling, combined with chemical probing, evolutionary covariation, and compensatory mutagenesis, to derive a biologically supported global secondary structure. To determine whether structurally distant regions could interact in the context of the full-length RNA, we engineered a structure-guided T1-ZD mutant that preserved the supported secondary structure while reducing alternative base-pairing possibilities. Full-length AlphaFold3 modeling of T1-ZD largely recapitulated the proposed architecture, while one predicted model revealed a long-distance interaction that was subsequently supported by compensatory mutagenesis analysis. These findings suggest that the 3′ terminus of sat-T1 may undergo conformational switching between alternative structural states. Together, our work establishes a biologically supported global RNA architecture for CMV sat-T1 and provides a structural framework for investigating the molecular basis of satRNA function.

**AUTHOR SUMMARY:** Satellite RNAs are small RNA molecules that depend on viruses for replication, yet they can strongly influence virus infection and disease symptoms in plants. Their biological activities are often determined by how the RNA folds into specific structures. However, understanding the overall structure of a long RNA remains challenging. Here, we established a biologically supported global structure for a satellite RNA associated with cucumber mosaic virus. Initial prediction of the complete RNA sequence using AlphaFold3 generated long-distance interactions that were not supported by biological evidence. Thus, we adopted the fragment-based strategy to generate an integrated global RNA model, which was supported by multiple experimental evidence. Finally, we engineered a mutant guided by the predicted structure and identified a biologically relevant long-distance interaction in the prediction models of the full-length mutant RNA. Our study provides a structural basis for understanding satellite RNA function and illustrates how experimentally supported RNA structures can guide artificial intelligence-assisted prediction of long RNA structures.

## INTRODUCTION

Satellite RNAs (satRNAs) are subviral agents that depend on their helper viruses for replication, encapsidation, and systemic movement, yet they can profoundly influence virus accumulation and disease development in infected plants [1–3]. Except for a small subset of larger satRNAs with coding capability, all satRNAs are non-coding RNA molecules, which achieve their biological functions predominantly through RNA determinants or structured elements. Establishing biologically supported RNA structural models is therefore essential for understanding the molecular mechanisms underlying satRNA biology.

Extensive studies of plant virus-associated satRNAs have demonstrated that RNA structural elements function as key regulatory modules throughout the infection cycle. In turnip crinkle virus (TCV)-associated sat-C, dynamic conformational switching within the highly structured 3′ region coordinates sat-C replication, template repair, and symptom modulation, providing one of the best-characterized examples of RNA structure-dependent regulation in plant subviral RNAs[4–7]. Another typical example is bamboo mosaic virus satRNA (satBaMV) that encodes a P20 protein required for systemic trafficking in plants [8]. In addition, satBaMV contains conserved structural elements in both untranslated regions, including a 5′-terminal apical hairpin required for replication competition with the helper virus and three 3′ stem-loops essential for efficient satRNA propagation [9–11]. In addition, studies of satRNAs have increasingly extended beyond individual structural motifs toward defining their global RNA architecture. For example, the global conformation of tobacco mosaic virus (TMV)-associated satRNA has been investigated using atomic force microscopy, cryo-electron microscopy, and SHAPE-guided computational modeling, but its biological relevance remains to be systematically validated [12–14]. To date, the only biologically supported global RNA structure reported for a plant satellite RNA is that of Cymbidium ringspot virus (CymRSV)-associated satRNA (sat-Cym), established by integrating chemical probing and extensive mutational analyses [15]. Cucumber mosaic virus (CMV)-associated satRNAs have long served as well-established model systems for investigating RNA structure-function relationships [16]. CMV satRNAs are small linear non-coding RNAs (approximately 330-405 nt) that share no significant sequence similarity with the genomic RNAs of their helper virus [17]. Importantly, more than 180 full-length CMV satRNA sequences are currently available in GenBank [18], providing an exceptional evolutionary resource for covariation-guided structural validation, an advantage not currently available for most other plant satRNAs. Despite their compact genomes, CMV satRNAs profoundly influence viral replication, symptom development, and transmission through RNA-mediated mechanisms [18–23]. For example, a conserved gamma-shaped RNA element contributes to suppression of helper-virus replication and symptom attenuation [18]. In addition, specific sequence determinants underlie host-specific pathogenicity, exemplified by sat-Y-derived small interfering RNAs targeting the mRNA of tobacco magnesium protoporphyrin chelatase subunit I to induce plant yellowing [21, 22]and three nucleotides in sat-D that are required for systemic necrosis in tomato [23]. Together, these studies demonstrate that diverse biological functions of CMV satRNA are encoded by discrete functional RNA elements. Secondary structure models of several CMV satRNAs have been proposed using thermodynamic prediction constrained by enzymatic or chemical probing, yet none has been systematically evaluated by biological assays [24–30].

Artificial intelligence (AI)-based approaches have recently demonstrated remarkable capabilities in RNA structure prediction [31–34]. In particular, AlphaFold3 employs a diffusion-based deep-learning architecture to predict RNA structures directly at the atomic-coordinate level, providing a powerful framework for modeling global RNA architecture from sequence information [34]. Here, we applied AlphaFold3 to model the global RNA structure of CMV satRNA T1 (sat-T1) and evaluated the predicted structures using SHAPE-guided structural probing, evolutionary covariation analysis, and compensatory mutagenesis coupled with biological assays. We further used a structure-guided RNA engineering strategy to enable full-length AlphaFold3 modeling of sat-T1 and to examine potential long-distance interactions between distant regions of the RNA. Together, these approaches allowed us to establish a biologically supported global RNA architecture of sat-T1 and provided insights into its conformational plasticity.

## MATERIAL AND METHODS

### Plasmid construction

The infectious clones pCB301-F109, pCB301-F209, and pCB301-F309, corresponding to Fny-CMV RNA1, RNA2, and RNA3, respectively, have been described previously [35]. The infectious clone pCB301-T1, carrying the sat-T1 sequence, was reported previously [18]. Mutant derivatives of sat-T1 were generated by site-directed mutagenesis of pCB301-T1, in which specific nucleotides were substituted using PCR-based mutagenesis as previously described [18, 36]. For in vitro transcription, the sat-T1 sequence was amplified by PCR using primers pmT7-T1F and pmT1R-5H2-M13, which introduced a T7 promoter at the 5′ end and the PEMV-derived 5H2 hairpin [37], followed by an M13 sequence at the 3′ end. The resulting PCR fragment was cloned into pUC18 between the *Bam*HI and *Sma*I restriction sites. All plasmid constructs were verified by Sanger sequencing. Primers mentioned here and blow are listed in S1Table.

### 3D structure modeling and structural analysis

3D structure prediction was performed using the online AlphaFold3 server [34]. The full-length sat-T1 RNA sequence, the 5′ and 3′ RNA fragments, and full-length mutant sequences were used as input for modeling. Predictions were carried out using AlphaFold3 with the seed parameter set to “auto”, and no ions or additional molecular partners were included in the modeling system. For each input sequence, five structural models were generated by AlphaFold3. To extract RNA secondary structure information from the predicted 3D coordinates, canonical base-pairing interactions, including potential pseudoknots, were analyzed using RNApdbee 2.0, in which FR3D was used to identify canonical base pairs with removal of isolated base pairs [38]. The resulting base-pairing patterns were subsequently used to generate RNA secondary structure diagrams using RNAcanvas [39].

### SHAPE probing

Selective 2′-hydroxyl acylation analyzed by primer extension (SHAPE) was performed to determine nucleotide flexibility across the sat-T1 RNA, according to the procedures described previously [18, 40]. For RNA preparation, sat-T1 transcripts carrying a 3′-terminal 5H2-M13 sequence (T1-5H2-M13) were generated by *in vitro* transcription using the HiScribe T7 High Yield RNA Synthesis Kit (New England Biolabs) according to the manufacturer’s instructions. Approximately 3 pmol of purified RNA was folded in 1×folding buffer (80 mM Tris-HCl pH 8.0, 160 mM NH_4_Cl, and 11 mM Mg(OAc)_2_) by heating at 65 °C followed by incubation at 37 °C. Folded RNA was divided into two equal aliquots and treated with either the SHAPE reagent *N*-methylisatoic anhydride (NMIA) or dimethyl sulfoxide (DMSO) as a negative control. After chemical modification RNA was recovered and subjected to primer extension reactions. Primer extension reactions were performed using fluorescently labeled primers complementary to the M13 sequence. NMIA- and DMSO-treated RNAs were reverse-transcribed using a FAM-labeled primer to generate SHAPE signals. In parallel, sequencing ladders were generated from untreated RNA templates using a PET-labeled primer in the presence of dideoxycytidine-triphosphate (ddCTP). The sequencing ladders were mixed with either NMIA or DMSO cDNA samples, followed by capillary electrophoresis. Fluorescence traces were processed and quantified using QuShape [41], which performs peak identification, signal alignment, and normalization of SHAPE reactivities. The resulting normalized reactivity values were mapped onto the sat-T1 secondary structure using RNAcanvas for visualization [39].

### Covariation analysis

CMV satRNAs are classified phylogenetically into two groups [24], which exhibit a highly divergent sequence in their central region [17, 19]. To accurately assess covariation of CMV satRNAs, we retrieved 97 full-length sequences of group I satRNAs from the GenBank database (Supplementary Data 1). Sequences were aligned using MAFFT and manually inspected and adjusted in AliView [42] to optimize alignment of conserved structural regions. Covariation analysis was performed using R-scape [43] to evaluate statistically significant covarying base pairs within the alignment based on the predicted secondary structure model. The default R-scape statistical framework was applied, and base pairs with an E-value ≤ 1.0 were considered indicative of significant covariation supporting the predicted pairing interactions.

### Agroinfiltration assays

Agrobacterium-mediated virus inoculation was performed as previously described [18]. All T-DNA plasmids were transformed into *Agrobacterium tumefaciens* GV3101 using the freeze-thaw method [44]. Agrobacterial cells carrying the infectious clones of Fny-CMV RNA1, RNA2, and RNA3, together with sat-T1 or its mutant derivatives, were cultured and mixed in equal proportions. Agrobacterium cells harboring 35S:P19, which expresses the RNA silencing suppressor P19, were included in the mixture. The bacterial suspensions were infiltrated into the sixth true leaves of approximately 4-week-old *Nicotiana benthamiana* plants. Plants were maintained in a controlled growth room under a 16 h light / 8 h dark photoperiod, with a light intensity of 150-200 μE m^-2^ s^-1^ and a temperature of 23-25 °C. At 3 days post-infiltration (dpi), infiltrated leaves from three independent plants were harvested and pooled for RNA extraction.

### Northern blot analysis

Northern blot assays were performed as previously described [18]. Briefly, total RNA was extracted from pooled infiltrated leaves using extraction buffer containing 0.05 M sodium acetate (pH 5.2), 0.01 M EDTA (pH 8.0), and 1% SDS. RNA samples were separated on a 1.5% denaturing agarose gel and transferred onto a positively charged nylon membrane (GE Healthcare). Digoxigenin (DIG)-labeled DNA oligonucleotide probes were used to detect CMV RNA3, sat-T1, and its mutant derivatives. The probe pbFR3(+):13-52 was used for detection of CMV RNA3. For sat-T1 mutants carrying substitutions within nucleotides 21-57, hybridization was performed using probe pbT1(+):292-318, whereas other mutants were detected using probe pbT1(+):21-57. The negative strands of sat-T1 and T1-ZD were detected using a DIG-labeled oligonucleotide identical to nucleotides 26-62 of sat-T1. Hybridization signals were detected using a DIG chemiluminescence detection kit (Roche) according to the manufacturer’s instructions. Band intensities were quantified using ImageJ, and relative RNA accumulation levels were calculated by normalizing to the wild-type sat-T1 signal. Probe sequences are listed in S2 Table.

### Reverse transcriptase-polymerase chain reaction (RT-PCR)

Genetic stability of sat-T1 mutants were assessed by RT-PCR and Sanger sequencing. In brief, total RNA (1 μg) extracted from infiltrated leaves was treated with DNase I (Promega) to remove residual plasmid DNA introduced during agroinfiltration. The DNase-treated RNA was then used as a template for first-strand cDNA synthesis using SuperScript III reverse transcriptase (Invitrogen) and the T1-specific reverse primer pmT1:R324-337. The resulting cDNA was amplified by PCR using primers pmT1:F1-22 and pmT1:R324-337. PCR products were separated by agarose gel electrophoresis, purified, and subjected to Sanger sequencing to assess the genetic stability of sat-T1 mutants during infection. Mutants carrying substitutions near the 5′ or 3′ termini were analyzed by 5′ or 3′ RACE, respectively, as described below.

### 5′ Rapid amplification of cDNA ends (5′ RACE)

5′ RACE was performed as previously described [45] with minor modifications. First-strand cDNA was synthesized from total RNA using SuperScript III reverse transcriptase (Invitrogen) and the gene-specific reverse primer pmT1:R324-337. The resulting cDNA products were separated on a 1% agarose gel, and bands of the expected size were excised and purified. Purified cDNA was tailed at the 3′ end with a poly(C) tract using terminal deoxynucleotidyl transferase (TdT; New England Biolabs) in the presence of dCTP, according to the manufacturer’s instructions. The tailed cDNA was subsequently amplified by PCR using Q5 High-Fidelity DNA Polymerase (New England Biolabs) with an oligo(dC)_12_ forward primer and T1-specific reverse primer pT1:R321-337. PCR products were gel-purified and subjected to Sanger sequencing using primer pT1:R321-337.

### 3′ RACE

3′ RACE was performed following established protocols [46]. T1 mutant RNAs were enriched by separating total RNA on a 1.5% denaturing agarose gel, followed by gel extraction using the “crush and soak” method [47]. Purified RNA was polyadenylated at the 3′ end using *E. coli* poly(A) polymerase (New England Biolabs) in the presence of ATP. First-strand cDNA synthesis was performed using SuperScript III reverse transcriptase (Invitrogen) with an oligo(dT)-adaptor primer [46]. The resulting cDNA was amplified by PCR using Q5 High-Fidelity DNA Polymerase with the T1-specific forward primer pmT1:F1-22 and the adaptor-specific reverse primer. PCR products were gel-purified and sequenced using primer pmT1:F1-22.

## RESULTS

### Experimental-guided refinement of an AlphaFold3-derived secondary structure model of CMV sat-T1

We initially modeled the full-length sequence of sat-T1 (337 nt) using AlphaFold3, which generated five structural models. In the secondary structures derived from these models (S1A Fig), Model 0 was used as a representative model to illustrate the initial full-length prediction in Fig 1A. The five models exhibited a highly similar overall architecture, indicating that the major structural features were consistently predicted. For example, the two apical hairpins (H1 and H2) and the internal pseudoknot within the previously characterized γ-shaped structure (γSS) (S1B Fig) [18] were consistently recovered in all five models (blue circles). In contrast, the sequences forming the basal stem (Bs) of γSS were not predicted to pair with each other; instead, the two regions indicated in red were separated and paired with distal sequences elsewhere in the RNA. Because the biological relevance of the Bs has not been established previously [18], these alternative base-pairing interactions raised uncertainty as to whether the predicted pairing pattern represented a biologically relevant conformation of sat-T1.

**Fig 1.**
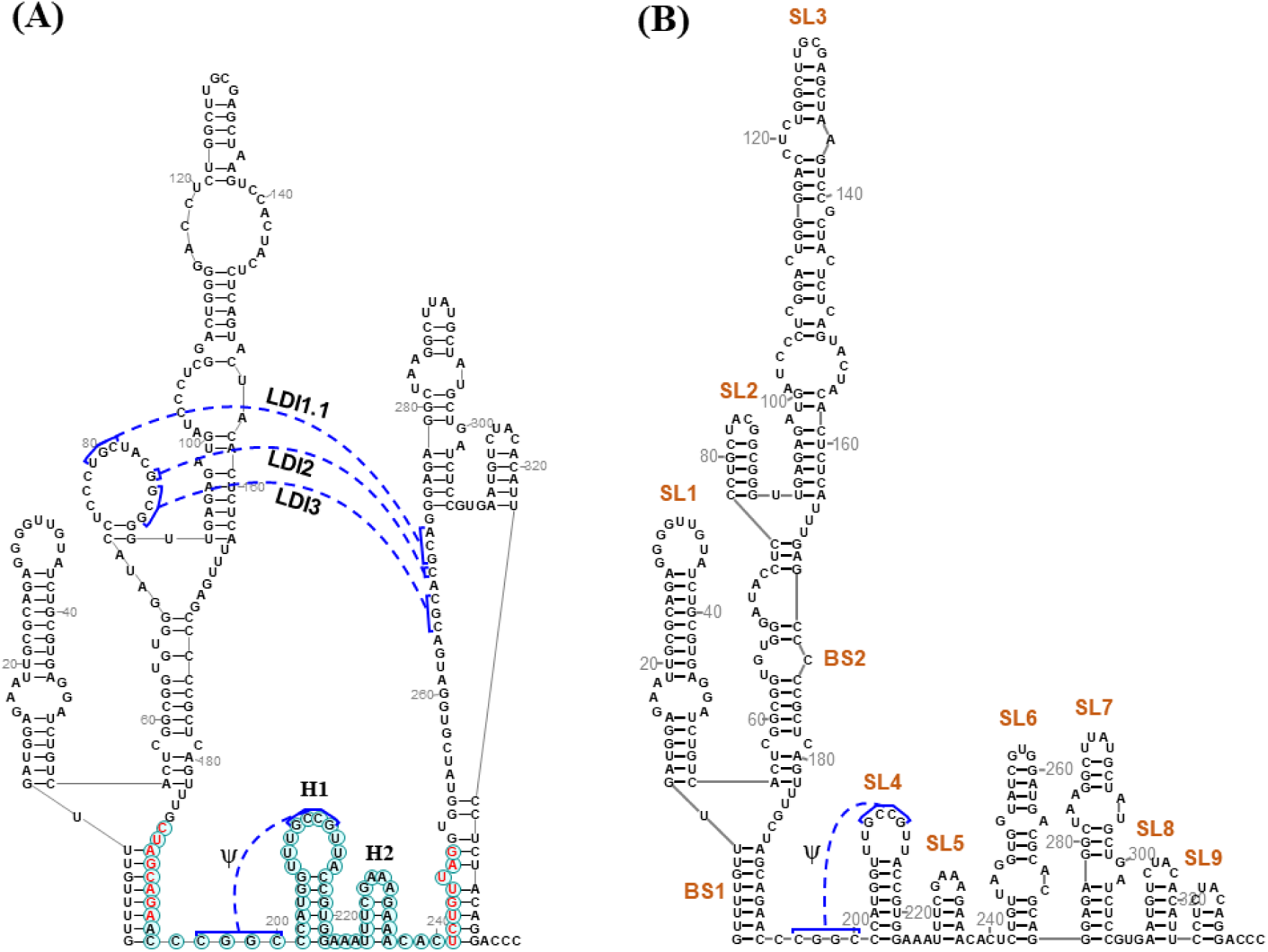
AlphaFold3-based modeling of the global secondary structure of sat-T1. (A) Representative full-length AlphaFold3 prediction of sat-T1 (Model 0). Green circles indicate the previously characterized H1, H2, and internal pseudoknot of the γ-shaped RNA structure (γSS), whereas the sequences colored red correspond to its basal stem (Bs). Predicted long-distance interactions (LDI1.1-LDI3) are indicated by blue, dotted lines. (B) Fragment-based AlphaFold3 modeling of two non-overlapping sat-T1 fragments (nt 1-195 and 196-337). Models 2 and 1 were selected as representative structures for the 1-195 and 196-337 fragments, respectively, and integrated with several local adjustments to generate the refined global secondary structure.

Next, we analyzed the long-distance interactions (LDIs) predicted across the five models. Three LDIs (LDI1-LDI3) were predicted in these models (S1A Fig). Specifically, LDI1 and LDI3 were consistently predicted across all five models, whereas LDI2 was detected in Models 0, 2, and 4. LDI1 comprised four alternative base-pairing patterns (LDI1.1-LDI1.4), which shared a common sequence interaction. To evaluate the biological relevance of the predicted LDIs, we selected LDI1.1 and LDI3 in Model 0 for mutational analysis (Fig 1A & S2A Fig). Disruption of either LDI1.1 or LDI3 interaction abolished sat-T1 accumulation (S2B Fig). However, compensatory mutations designed to restore the predicted base pairing failed to rescue sat-T1 accumulation. Thus, the lack of phenotypic rescue by compensatory mutations did not support these predicted interactions as biologically functional base-pairing interactions. These results prompted us to refine the modeling strategy by reducing the sequence context available for long-distance pairing.

To reduce the influence of unsupported long-distance interactions, we next divided sat-T1 into two non-overlapping fragments, comprising nucleotides 1-195 and 196-337, and modeled each fragment separately using AlphaFold3. The five predictions for the 1-195 fragment showed highly similar global architectures, characterized by a nested, branched RNA structure (S3 Fig). Notably, the sequences corresponding to the 5′ regions in LDI1.1 and LDI3 consistently formed a local stem-loop structure spanning nucleotides 77-90. Similarly, four of the five predictions for the 196-337 fragment (Models 0-3) exhibited nearly identical architecture, with five consecutive stem loops arranged in the same configuration (S4 Fig). The apical hairpins of γSS were retained in all five predictions, whereas the internal pseudoknot was predicted only in Model 1.

Based on the consistency among the fragment-derived structures and our prior structural knowledge of sat-T1, we selected Model 2 of the 1-195 fragment and Model 1 of the 196-337 fragment as representative structures and integrated them to refine a global secondary structure model, with several local adjustments (Fig 1B). These adjustments included refinement of the base-pairing pattern within SL3, folding of nucleotides 325-333 into a small terminal stem-loop (SL9), and disruption of the ^284^A:U^295^ base pair. The resulting model featured a nested branched architecture in the 5′ half, with SL2 and SL3 branching from the upper basal stem (BS2) of the primary branch, followed by six consecutive stem-loop elements (SL4-SL9) in the 3′ half, with a SL4 participating in pseudoknot formation. This refined model incorporated structural features supported by the fragment predictions while avoiding the unsupported long-distance interactions identified in the initial full-length models. It therefore provided a defined structural hypothesis for subsequent evaluation by SHAPE probing, evolutionary covariation analysis, and compensatory mutagenesis.

### Covariation and SHAPE probing validate the refined sat-T1 structure

To evaluate the refined secondary structure model, we performed covariation analysis using the R-scape algorithm based on a multiple-sequence alignment of 97 full-length CMV satRNA sequences (Fig 2). Overall, CMV satRNAs exhibit high sequence conservation (colored red), with sequence variability largely confined to loop or bulge regions, such as the bulge spanning nt 96-99 and the loop between nt 287 and 293. In the refined model, eight covarying base pairs were identified (boxed in green). Of these, three occur within SL3, providing strong support for the reliability of this structural element. The remaining five covarying pairs are distributed among four additional stem-loops (SL4, SL6, SL7, and SL8), further supporting the local structural features predicted by AlphaFold3.

**Fig 2.**
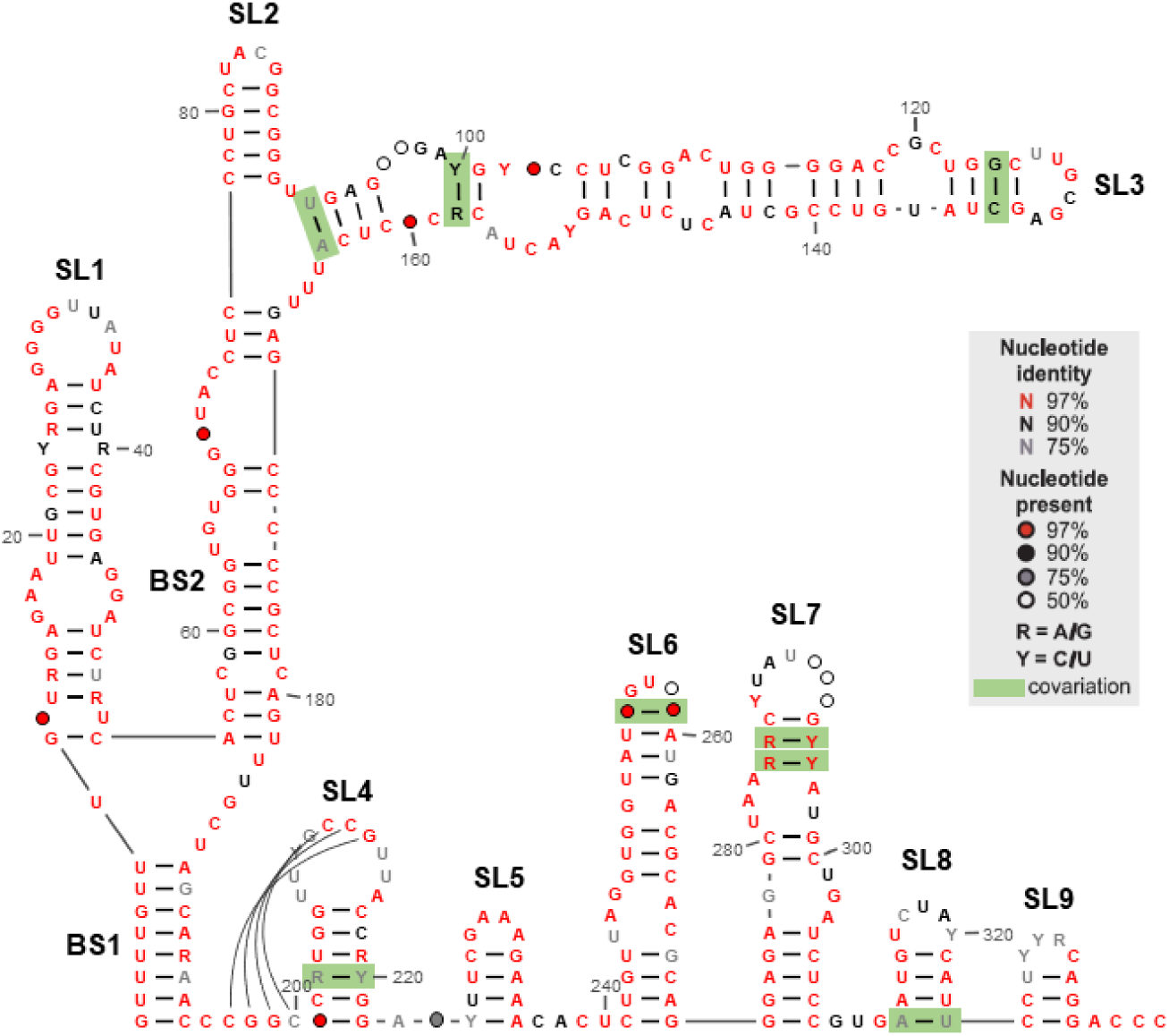
Covariation analysis supports the predicted sat-T1 secondary structure. A multiple sequence alignment of 97 CMV satellite RNAs was analyzed using R-scape. Covarying base pairs are highlighted in green boxes and mapped onto the predicted secondary structure. Nucleotide conservation is indicated by color coding or circle annotations.

We next assessed the structure model using SHAPE assays (Fig 3). A total of 84 nucleotides exhibited detectable reactivity, including 56 highly reactive (colored brown) and 28 moderately reactive bases (colored blue). Most reactive nucleotides were located in loops and bulges or adjacent to base-paired regions, consistent with their expected structural accessibility. Notably, some paired bases also displayed strong reactivity, whereas some unpaired bases showed little or no reactivity. These discrepancies likely reflect local conformational dynamics or alternative structural states of sat-T1 in solution. Overall, the covariation and SHAPE probing data provided independent support for the refined secondary structure model.

**Fig 3.**
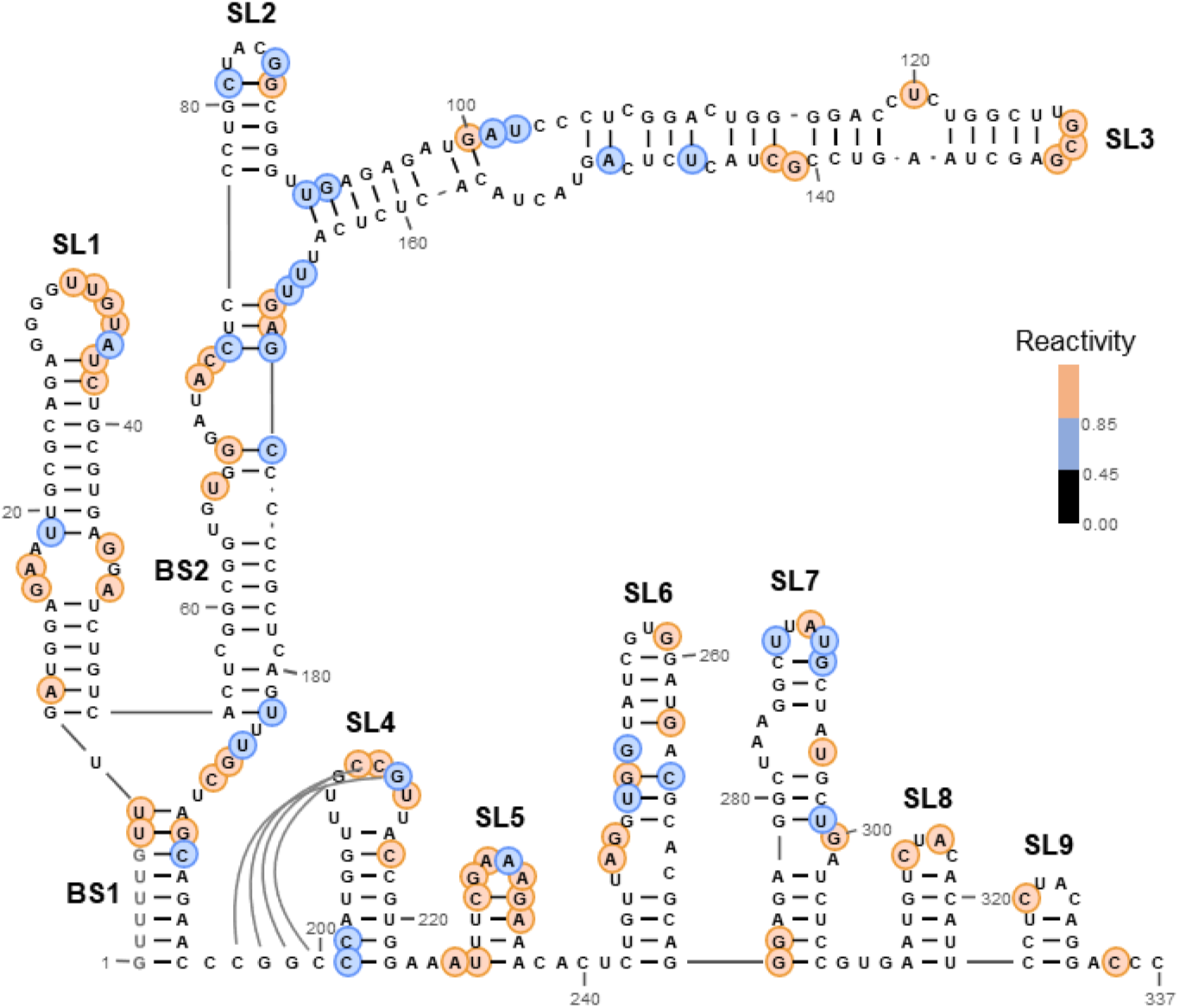
SHAPE probing validates the global folding of sat-T1 RNA. *In vitro*-transcribed sat-T1 RNA, containing a 3′ terminal extension derived from pea enation mosaic virus 5H2 and M13 sequences, was subjected to SHAPE (Selective 2′-Hydroxyl Acylation analyzed by Primer Extension) analysis to assess nucleotide flexibility. Reactivity values are mapped onto the refined secondary structure. Highly (>0.85) and moderately (0.45-0.85) reactive nucleotides are shown in brown and blue, respectively. The first six nucleotides at the 5′ terminus produced unreliable fluorescence signals and were therefore excluded from reactivity calculations and masked in grey.

### Functional validation of the predicted structural model

In the predicted model, the biological significance of SL4, including its participation in the pseudoknot (Pk1) formation, as well as SL5, has been documented previously [18]. To evaluate the biological relevance of the remaining structural elements, substitutions were introduced into helical regions and terminal loops, as described below. Mutants designated with the suffixes “A” or “B” contain disruptive substitutions in helical regions, whereas those ending in “C” carry compensatory mutations. For terminal loops, loop sequences were replaced with complementary nucleotides. All mutants were tested in *N. benthamiana* plants with co-infection of CMV via agroinfiltration. Genetic stability of all mutants was confirmed by RT-PCR, 5′ RACE, or 3′ RACE, followed by sequencing.

### Nested branching structure at 5′ region

The nested branching structure is composed of two basal stems (BS1 and BS2) and three stem-loops (SL1, SL2, and SL3) (Fig 4). The base pairs selected for BS1 mutagenesis warrant special attention because its 5′ complementary sequence (GUUUUGUU) locates at the exact 5′ terminus and is highly conserved. To generate the disruptive mutant (B1A), nucleotides uridine at position 5 (^5^U) and guanine at position 6 (^6^G) were substituted with adenine (A) and U, respectively, yielding a sequence identical to the 5′ terminus of CMV RNA3. Disruption of the BS1 stem (B1A and B1B) markedly reduced sat-T1 accumulation (> 75%), whereas the corresponding compensatory mutation (B1C) restored accumulation. Similarly, most helical regions (B2.1-B2.4) within BS2 were required for sat-T1 accumulation, as their disruptive mutants were functionally defective, with the exception of B2.4B, which accumulated at a level comparable to wild type. For SL1, disruption of either lower helix (S1.1) or upper helix (S1.2), as well as substitution of the terminal loop sequence, were detrimental to sat-T1 survival, while both compensatory mutants (S1.1C, S1.2C) restored accumulation level comparable to wild type. These results indicate an essential role for this element. SL2 is a small hairpin branching from BS2; both its helical structure and terminal loop were functionally important, with the former being essential for its viability. SL3 is the second branch containing 7 helical regions (S3.1-S3.7) separated by internal loops. Disruption of all SL3 helices impaired sat-T1 accumulation, with the lower helix (S3.1) and the apical helices (S3.6 and S3.7) being particularly critical. In contrast, disruption of the internal helix S3.5 was tolerated. Substitution of the terminal loop also affected accumulation but was less detrimental than disruption of the apical helices. Collectively, these results demonstrate that all structural elements forming the nested branching architecture are biologically significant.

**Fig 4.**
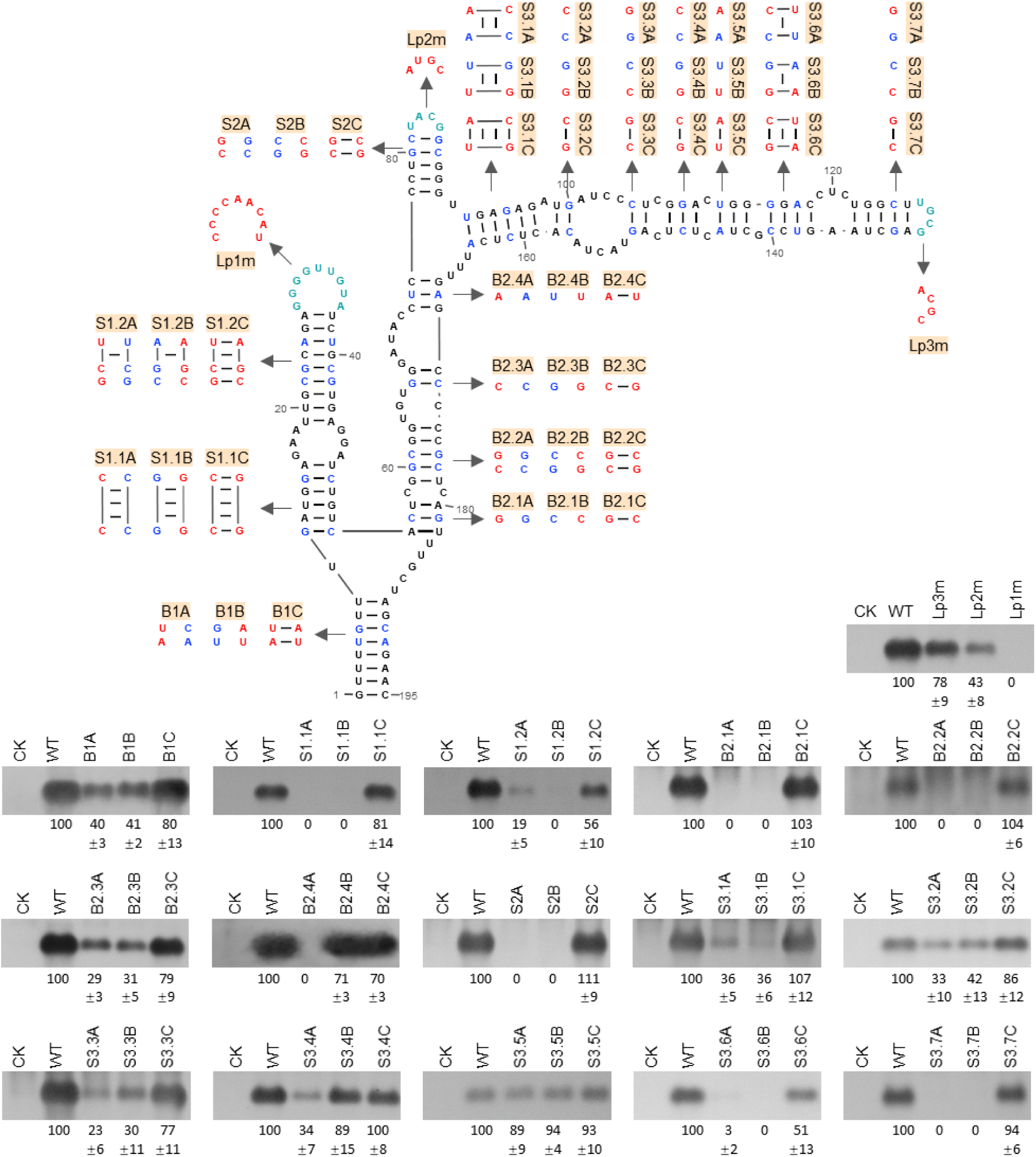
Functional validation of the 5′ nested branching structure. Base pairs and terminal loops selected for mutational analysis are highlighted in blue and green, respectively. All mutants were co-inoculated with CMV into Nicotiana benthamiana plants via agroinfiltration. At 3 days post-infiltration (dpi), RNA accumulation of sat-T1 and its mutants in infiltrated leaves was analyzed by Northern blotting. Representative results are shown for each set of mutants. Relative accumulation levels represent the mean ± s.e.m. from three independent biological replicates.

### SL6

SL6 comprises three helical regions (S6.1-S6.3) capped by a three-nucleotide terminal loop (Fig 5). The biological relevance of these helices was assessed by compensatory mutation analysis (Fig 5A). Disruptive mutations (S6.1A, S6.1B) in the lower helix severely impaired sat-T1 accumulation, whereas the compensatory mutation (S6.1C) restored function, indicating the importance of this helical structure. For the upper helix (S6.3), the top base pair (^256^C:G^260^) covaries in CMV satRNAs (Fig 2). However, both disruptive mutations (S6.3A, S6.3B) and the corresponding compensatory mutation (S6.3C) were detrimental. Similar outcomes were observed for mutations targeting the neighboring base pair ^255^U:A^261^ (S6.3D-F) and the base pair ^249^G:C^267^ (S6.2A-C) in the middle helix. These findings suggest that, in addition to helical integrity, specific nucleotide identities within SL6 are important for function.

**Fig 5.**
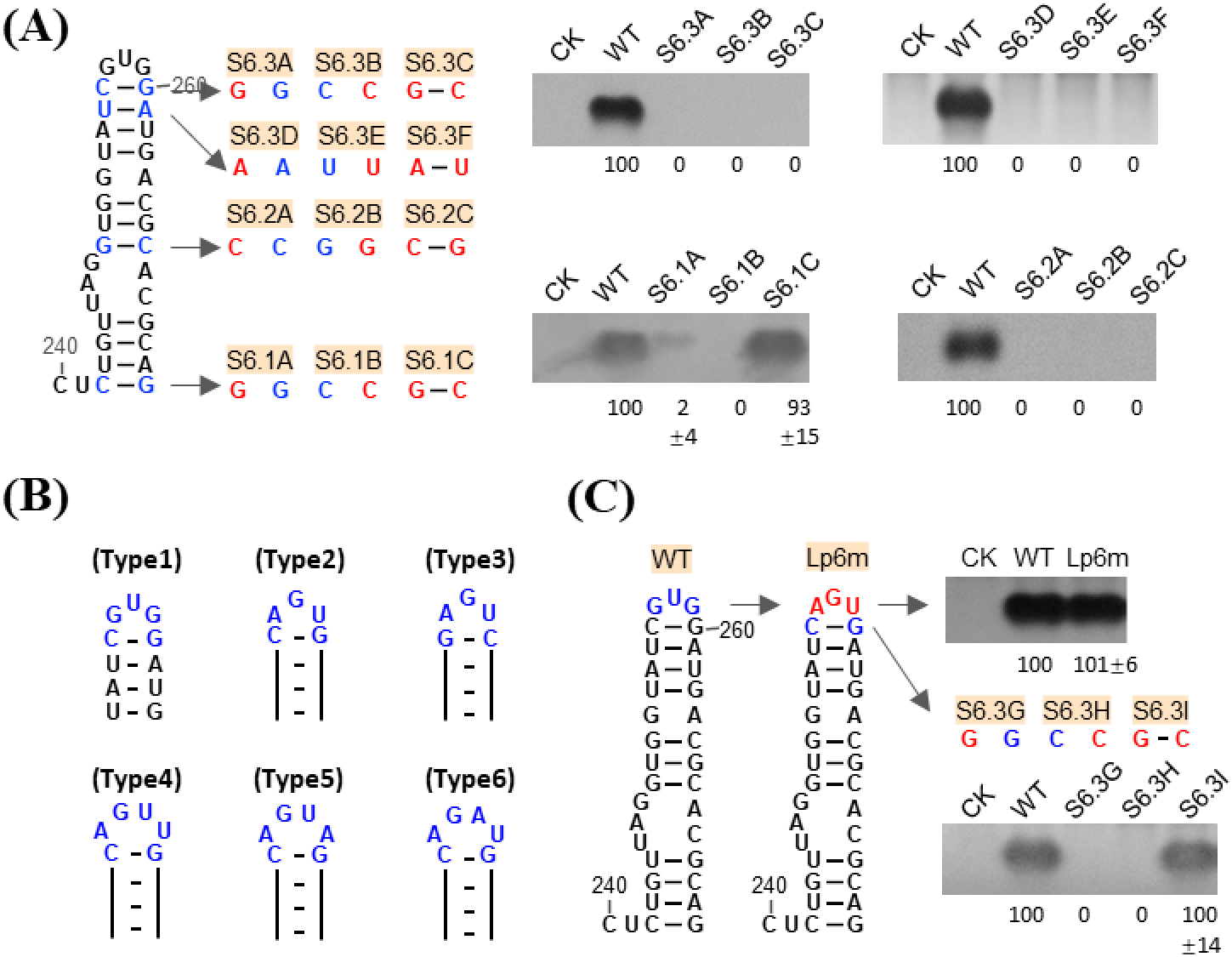
Functional analysis of SL6. (A) Compensatory mutation analysis of SL6 helices. Base pairs selected for mutational analysis are highlighted in green, and introduced mutations are shown in red. Mutants were tested in Nicotiana benthamiana as described in Fig 4. (B) Sequence variation of the SL6 terminal loop among CMV satellite RNAs. Variants are grouped into six types (Type 1-6). Covariation of the 256:260 base pair is observed specifically in sequences with an AGU loop (Type 2 and Type 3). (C) Functional rescue of compensatory mutations in a loop-modified background. Substitution of the loop sequence GUG with AGU generated the mutant Lp6m (red). In this background, the ^256^C:G^260^ base pair (blue) was subjected to compensatory mutation analysis. Mutants were tested as described in Fig 4.

Although the behavior of the compensatory mutant S6.3C appeared inconsistent with the observed covariation, sequence analysis of 125 CMV satellite RNAs revealed that covariation of ^256^C:G^260^ to G:C occurs only when the terminal loop sequence is AUG (Types 2 and 3; Fig 5B). Substitution of the SL6 terminal loop from GUG to AGU (Lp6m) was functionally acceptable (Fig 5C). In the Lp6m background, the compensatory mutation (6.3I) of ^256^C:G^260^ successfully rescued the defects of the corresponding disruptive mutants (S6.3G and S6.3H). These results indicate that the upper helical structure of SL6 is functionally relevant in a sequence-dependent manner.

### SL7

SL7 contains three helical regions (S7.1-7.3), each with a single G:C pair that was individually subjected to compensatory mutation analysis (Fig 6). Introducing mismatches into any of these helices markedly reduced sat-T1 accumulation, with the C-C mismatch (S7.3A) in the upper helix having the most severe effect. In all cases, the corresponding compensatory mutations (S7.1C, S7.2C, S7.3C) fully restored accumulation. Similarly, substitution of the terminal loop sequence caused moderate effect on its accumulation. These results indicate that both the stem and the terminal loop of SL7 play an important role for sat-T1 accumulation.

**Fig 6.**
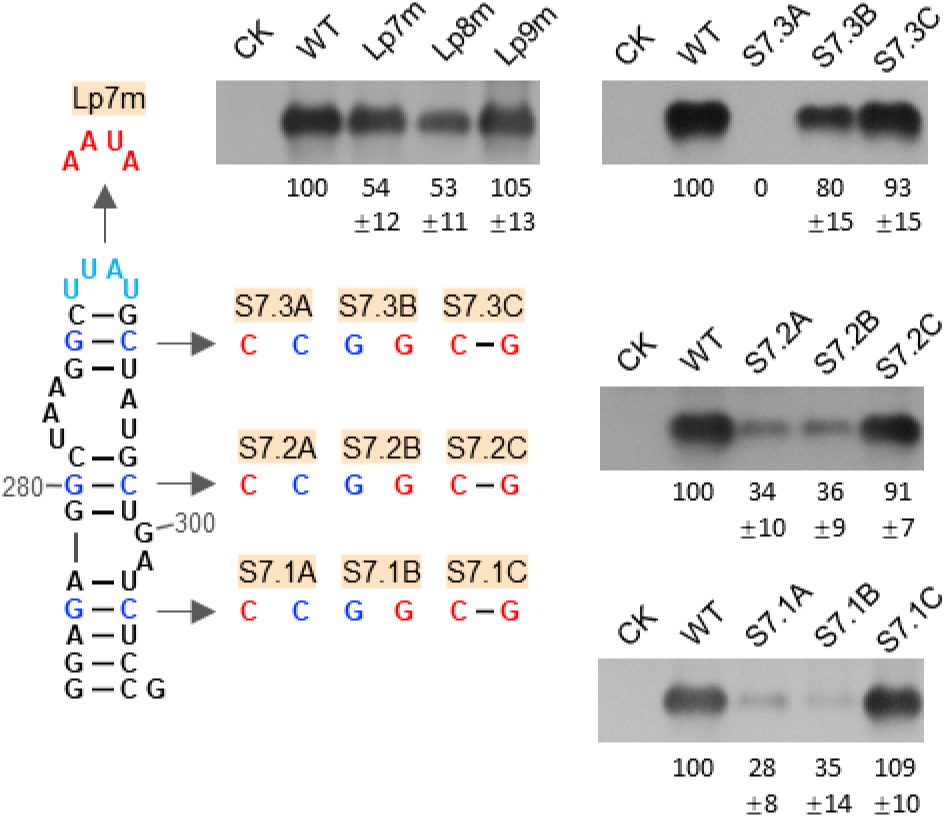
Functional analysis of SL7. Secondary structure of SL7 with disruptive and compensatory mutations indicated. Mutants were analyzed in Nicotiana benthamiana as described in Fig 4.

### SL8 & SL9

The 3′ terminal region of sat-T1 folds into two small stem-loops (SL8, SL9), followed by four unpaired nucleotides (ACCC) at the exact 3′ end. Disruptive mutations in either SL8 (S8A, S8B) or SL9 (S9A, S9B) reduced sat-T1 accumulation by more than 67%, whereas the corresponding compensatory mutations (S8C and S9C) substantially restored accumulation (Fig 7A). Substitution of the SL8 terminal loop (Lp8m) caused a moderate reduction in accumulation, whereas substitution of the SL9 terminal loop (Lp9m) had no detectable effect (Fig 6). These results demonstrate the biological importance of SL8 and SL9, depending primarily on their helical structures.

**Fig 7.**
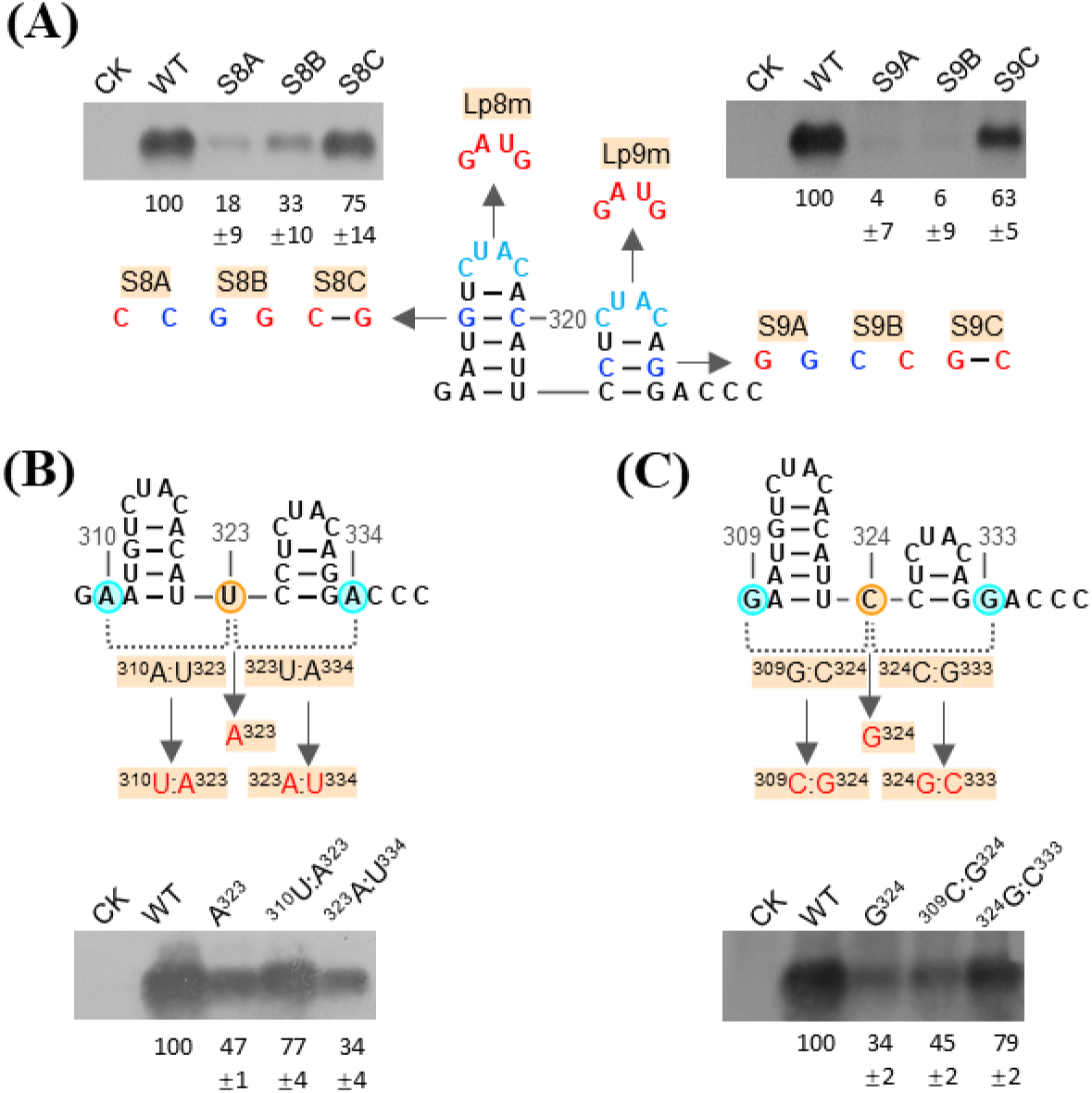
Structural and functional characterization of SL8 and SL9. (A) Functional analysis of SL8 and SL9. Base pairs and terminal loops selected for mutational analysis are highlighted in blue and green, respectively. Mutants were tested as described in Fig 4. (B) Analysis of alternative base pairing involving U^323^. U^323^ is predicted to pair with either A^310^ or A^334^ (dotted lines). U^323^ was mutated alone or in combination with mutations at position 310 or 334 (red). Mutants were analyzed in *Nicotiana benthamiana*, and RNA accumulation was assessed by Northern blotting. (C) Analysis of alternative base pairing involving C^324^. C^324^ is predicted to pair with either G^309^ or G^333^ (dotted lines). Mutational analyses were performed as described for U^323^, and RNA accumulation was examined as above.

Within these stem-loops, U^323^ could potentially pair with A^310^ or A^334^ (Fig 7B). To determine biologically relevant interaction, U^323^ was substituted with A^323^, disrupting both potential pairings, which reduced sat-T1 accumulation by 53%. This reduction was partially restored by the compensatory mutation at position 310 (^310^U:A^323^), but not by the mutation at position 334 (^323^A:U^334^) (Fig 7B), demonstrating that the ^310^A:U^323^ pair contributes to SL8 but not SL9. Similarly, C^324^ could potentially pair with G^309^ or G^333^ (Fig 7C). Substitution of C^324^ with G (G^324^) reduced sat-T1 accumulation by 64%, which was rescued substantially when this mutation was combined with the mutation C^333^ (^324^G:C^333^), but not with the mutation C^309^ (^309^C:G^324^). These results indicate that the ^324^C:G^333^ base pair contributes to the functional SL9 structure (Fig 7C). Collectively, these findings support the predicted structures of SL8 and SL9 and underscore their essential roles in sat-T1 activity.

### Modeling the global sat-T1 structure in a full-length RNA context using a T1 mutant

Fragment-based modeling cannot determine whether structurally distant regions form biologically relevant interactions in the context of the full-length RNA. We therefore engineered T1-ZD, a structure-guided sat-T1 mutant designed to reinforce the experimentally supported base-pairing interactions and disfavor alternative local folds (Fig 8A). T1-ZD incorporated six sets of compensatory mutations targeting alternative structures identified during the analysis, including S2C to disfavor the predicted LDI1 and LDI3 interactions, S3.4C and S3.6F to correct local mispairing within SL3, 324G:C333 to favor the experimentally supported SL9 structure, and B2.3C to prevent a local G-register shift. Each structural mutation was individually supported by biological assays (Figs 4 & 7; S5 Fig). T1-ZD retained biological activity in planta, although its positive- and negative-strand RNA accumulation was reduced by approximately 48% and 59%, respectively, relative to parental sat-T1 (Fig 8B), indicating that the engineered RNA preserved the essential structural features required for sat-T1 propagation.

**Fig 8.**
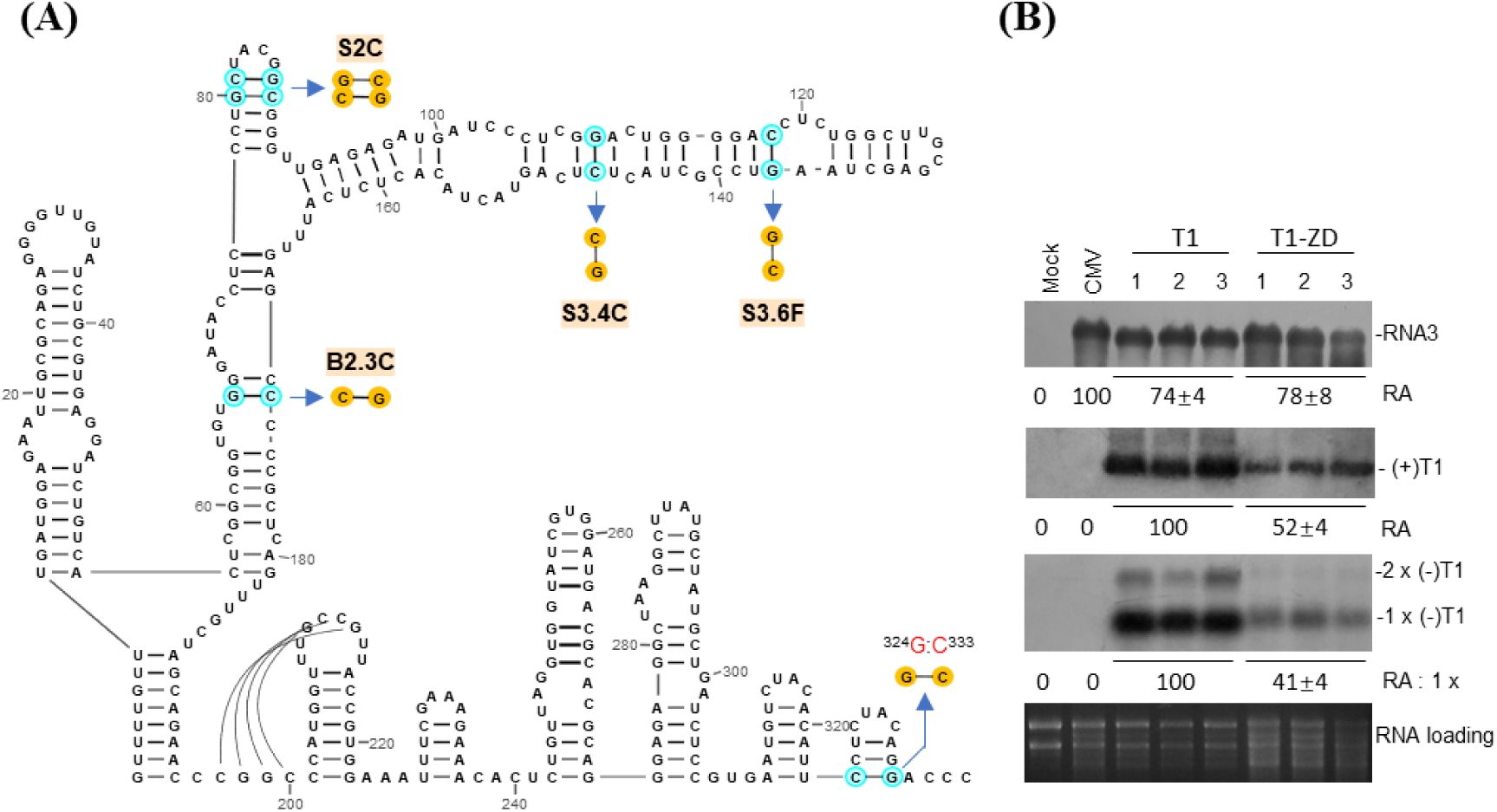
Design and validation of the structure-guided mutant T1-ZD. (A) Schematic of introduced compensatory mutations. Six pairs of compensatory mutations were introduced into sat-T1 to generate T1-ZD. (B) Biological activity test of T1-ZD. T1-ZD was co-inoculated with CMV into *Nicotiana benthamiana* via agroinfiltration. At 3 dpi, northern blot analysis was performed to examine the accumulation of positive- and negative-strand sat-T1 and T1-ZD RNAs, as well as CMV RNA3. Relative accumulation levels represent the mean ± s.e.m. from three biological replicates. Ethidium bromide-stained rRNAs serve as loading controls.

We next used the full-length T1-ZD sequence as input for AlphaFold3 and examined the five predicted models for two features: recapitulation of the experimentally supported secondary structure and formation of additional long-distance interactions. Models 1, 3, and 4 fully recapitulated the supported secondary structure, whereas Models 0 and 2 showed the same overall architecture with only limited local deviations (S6 Fig & Fig 9A). Thus, the experimentally supported architecture could be largely reproduced from the full-length T1-ZD sequence. Notably, Model 2 predicted a potential long-distance interaction between the 3′-terminal ACC sequence (nt 334-336) and the GGU sequence (nt 30-32) in the SL1 loop (Fig 9A). Formation of this interaction was accompanied by disruption of the base pairing required for SL9, leaving the corresponding SL9-forming sequence unpaired (colored blue). Model 2 therefore revealed an alternative conformation in which the 3′ terminus engages the 5′ region of sat-T1.

**Fig 9.**
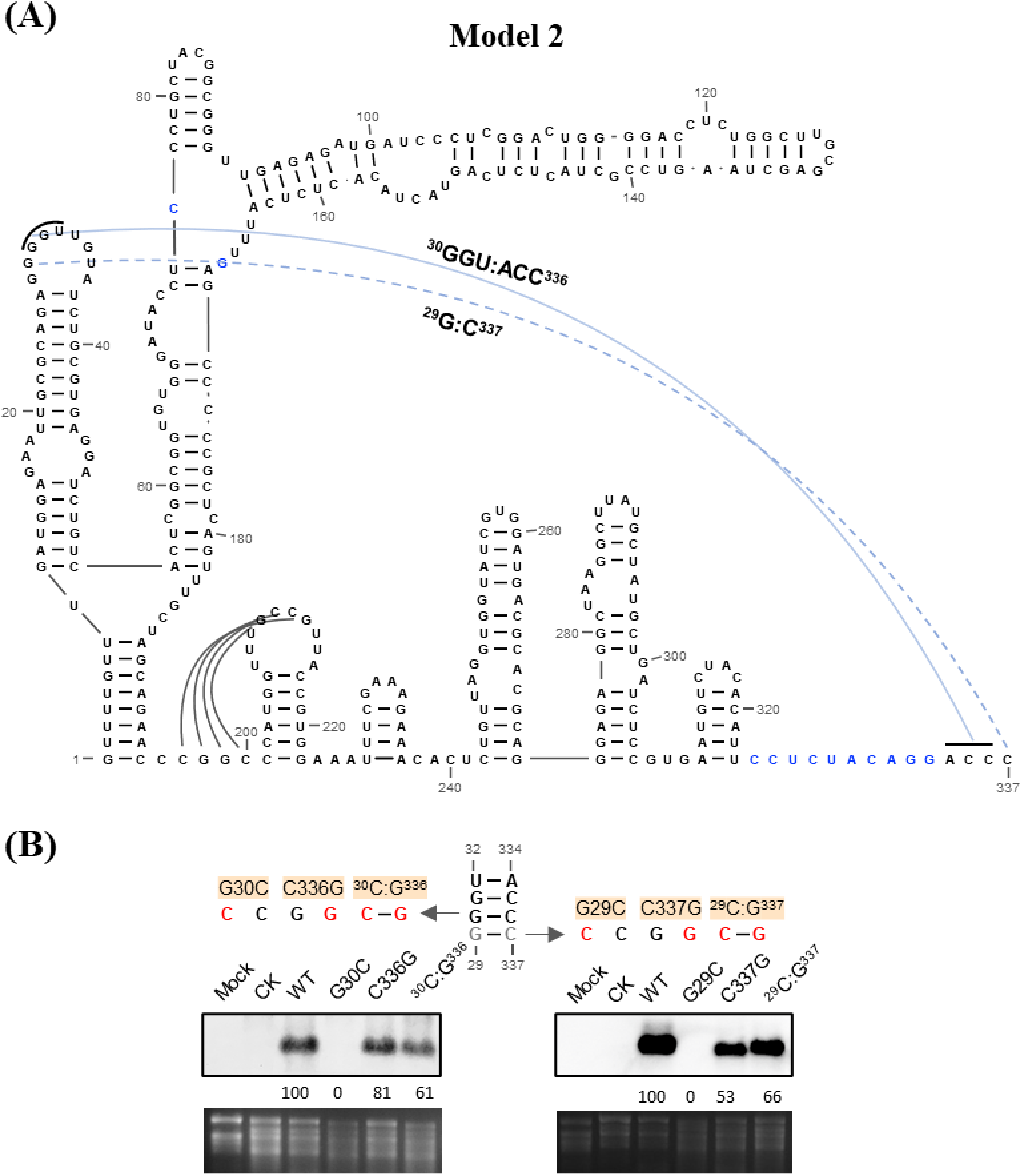
Full-length AlphaFold3 modeling of T1-ZD identifies a biologically supported long-distance interaction. (A) A representative model (Model 2) of T1-ZD. The predicted secondary structure largely recapitulates the experimentally supported sat-T1 architecture. The 3′-terminal ACC sequence (nt 334-336) was predicted to base-pair with the GGU sequence (nt 30–32) in the SL1 loop, accompanied by disruption of the SL9-forming base pairs (blue). (B) Biological validation of the predicted long-distance interaction. Disruptive and compensatory mutations targeting the predicted ^30^G:C^336^ and ^29^G:C^337^ base pairs were introduced into sat-T1, and RNA accumulation was analyzed by Northern blotting in *Nicotiana benthamiana*.

We then asked whether this predicted long-distance interaction was biologically relevant. Substitution of G^30^ with C (G30C) severely impaired sat-T1 accumulation to an undetectable level, as reported previously (He et al., 2019), whereas the reciprocal mutation C336G had limited effect (Fig 9B). Combining the two mutations restored sat-T1 accumulation to 61% of the wild-type level. Although G^29^ was not predicted to pair with C^337^ in Model 2, testing the adjacent ^29^G:C^337^ pair produced a similar compensatory effect (Fig 9B). Together, these results provide biological support for the predicted long-distance interaction and further support a potential base-pairing interaction between G^29^ and C^337^.

## DISCUSSION

In this study, we established a global secondary structure of CMV sat-T1 using a fragment-based modeling strategy supported by biochemical, evolutionary, and functional evidence. Importantly, extending this framework to a structure-guided full-length T1-ZD sequence enabled AlphaFold3 to reproduce the proposed architecture while revealing a long-distance interaction that was not captured by the fragment-based model. Functional validation of this interaction further suggests that the 3′ terminus of sat-T1 may undergo conformational switching between alternative structural states. Thus, the global architecture of sat-T1 provides a structural framework for understanding how RNA architecture contributes to CMV satRNAs functions. Moreover, we developed an iterative strategy that combines fragment-based structural AI modeling with structure-guided full-length prediction, offering a practical approach to resolving the global architecture of long RNAs.

The sat-T1 model represents the first biologically supported global secondary structure model for CMV satRNA. Comparison of the sat-T1 model with previously proposed structures of evolutionarily related CMV satRNAs reveals both conserved and distinctive structural features. Several local elements of sat-T1 are also predicted in other CMV satRNAs. For example, SL1 is present in the proposed structures of satRNAs IX, D4, and Y [25, 27, 28, 48], whereas the BS1 element involving the extreme 5′ region has been reported only in the *in vitro* model of satRNA D4 [25]. SL8 is predicted in satRNAs B2, B3, G, and WL1[29], while SL9 is present in the proposed structures of satRNAs Q, D4, and TA-Tb [24, 26, 30]. It is worth mentioning that SL8 and SL9 have not been predicted together in the previously reported models. Moreover, several prominent features of sat-T1 model, including BS2, SL3, and SL6, have not been predicted previously. The biological relevance of SL3 and SL6 is further supported by evolutionary covariation, providing independent evidence for these structural elements. In addition, our model contains two biologically supported tertiary structural interactions: one pseudoknot and one LDI, neither of which has been reported in previously proposed satRNA structures. Together, these comparisons indicate that sat-T1 shares several structural elements with other CMV satRNAs while also exhibiting distinctive higher-order features.

An important implication of the global sat-T1 structure is that it provides a structural context for interpreting functional RNA elements previously characterized largely at the sequence level. For the γSS element, previous compensatory mutagenesis of its basal stem did not reveal a significant effect on sat-T1 accumulation, leaving the biological relevance of this predicted structural feature unresolved [18]. In the global structure established here, however, the sequences forming this basal stem are engaged in additional base-pairing interactions that were independently supported by our mutational analyses, suggesting that the structural context of this region is more complex than previously appreciated. Similarly, the host-specific pathogenicity of CMV satRNAs has been linked to defined sequence determinants. A related structural insight emerges from the necrogenic satRNAs, such as sat-D, in which three nucleotides within the apical region of SL7 have been shown to be essential for systemic necrosis in tomato [49]. Our model supports the localization of these determinants within SL7, while predicting different base-pairing relationships in the middle and basal helices of the stem compared with previously proposed structures [25]. These structural refinements provide a more complete view of SL7 and may help guide future investigation of how RNA architecture contributes to sat-D-induced tomato necrosis. Thus, the global structure does not by itself establish the molecular mechanisms underlying these pathogenicity determinants, but it places previously characterized functional sequences within a defined structural framework that can be experimentally tested.

A notable feature of the sat-T1 architecture is the structural plasticity of its 3′-terminal region. Our data support two alternative conformational states: one in which SL9 is formed and the 3′ terminus remains unpaired (Fig 1), and another in which SL9 is disrupted and the 3′-terminal ACC sequence engages in long-distance base pairing with the SL1 loop (Fig 9). Importantly, both structural states are functionally relevant, as disruption of SL9 impairs sat-T1 accumulation, whereas compensatory restoration of the predicted 3′-terminal interactions partially restores accumulation of corresponding mutants. These findings suggest that the 3′ terminus of sat-T1 may function as a conformational switch rather than adopting a single static structure. Similar alternative 3′-terminal conformations have been reported in tomato bushy stunt virus (TBSV) and cognate satellite RNAs (sat-B10), in which switching between open and closed states is linked to RNA replication and 3′-end accessibility [15, 50, 51]. In addition, the 3′ terminus of satRNAs associated with Cymbidium ringspot virus (CymRSV) or TCV also adopt dynamic conformations [4–7, 15], further supporting the idea that structural plasticity at the 3′ end may be a recurring feature of satRNA architecture.

The T1-ZD experiment further raises the possibility that structured-guided RNA sequence engineering could be used to guide AI-based structure prediction. Although this strategy does not directly impose structural constraints during model generation, it provides a proof of concept that experimentally supported structural information can be encoded into sequence space to bias the structural solutions sampled by an AI model. This concept is consistent with recent advances in experiment-guided AlphaFold3 modeling to generate protein structural ensembles consistent with experimentally observed conformations and dynamics[52]. For RNA, this concept may be particularly valuable for modeling RNA-protein complexes, where the biological relevance of RNA conformation and the geometry of protein-binding sites are closely coupled. In this regard, the structure-guided T1-ZD sequence may provide a useful experimental platform for testing whether biologically supported RNA architectures can improve AI-based modeling of interactions between CMV satRNA and viral or host proteins.

## ACKNOWLEDGEMENTS

We would like to thank our colleagues Liu Leyu and Lai Jialiang for their assistance with the experiments.

## DATA AVAILABILITY

All relevant data are within the manuscript and its Supporting Information files

